# Gene flow mediates the role of sex chromosome meiotic drive during complex speciation

**DOI:** 10.1101/024711

**Authors:** Colin D. Meiklejohn, Emily L. Landeen, Kathleen E. Gordon, Thomas Rzatkiewicz, Sarah B. Kingan, Anthony J. Geneva, Jeffrey P. Vedanayagam, Christina A. Muirhead, Daniel Garrigan, David L. Stern, Daven C. Presgraves

**Affiliations:** School of Biological Sciences, University of Nebraska, Lincoln, Nebraska, 68588 USA; Department of Biology, University of Rochester, Rochester, New York, 14627, USA; Janelia Research Campus, Howard Hughes Medical Institute, 19700 Helix Drive, Ashburn, Virginia, 20147, USA; Department of Integrative Biology, University of California, Berkeley, California, 94720, USA; Department of Molecular Biology and Genetics, Field of Genetics, Genomics, and Development, Cornell University, Ithaca, New York, 14853, USA; Pacific Biosciences of California, Menlo Park, CA USA; Department of Organismic and Evolutionary Biology, Harvard University, Cambridge, MA, 02138, USA; Department of Developmental Biology, Sloan-Kettering Institute, New York, NY 10065, USA; AncestryDNA, 153 Townsend Street, San Francisco, CA 94107 USA

## Abstract

During speciation, sex chromosomes often accumulate interspecific genetic incompatibilities faster than the rest of the genome. The drive theory posits that sex chromosomes are susceptible to recurrent bouts of meiotic drive and suppression, causing the evolutionary build-up of divergent cryptic sex-linked drive systems and, incidentally, genetic incompatibilities. To assess the role of drive during speciation, we combine high-resolution genetic mapping of X-linked hybrid male sterility with population genomics analyses of divergence and recent gene flow between the fruitfly species, *Drosophila mauritiana* and *D. simulans*. Our findings reveal a high density of genetic incompatibilities and a corresponding dearth of gene flow on the X chromosome. Surprisingly, we find that, rather than contributing to interspecific divergence, a known drive element has recently migrated between species, caused a strong reduction in local divergence, and undermined the evolution of hybrid sterility. Gene flow can therefore mediate the effects of selfish genetic elements during speciation.

## INTRODUCTION

Speciation involves the evolution of reproductive incompatibilities between diverging populations, including prezygotic incompatibilities that prevent the formation of hybrids and postzygotic incompatibilities that render hybrids sterile or inviable. Two patterns characterizing speciation implicate a special role for sex chromosomes in the evolution of postzygotic incompatibilities: Haldane’s rule, the observation that hybrids of the heterogametic sex preferentially suffer sterility and inviability (Haldane, 1922, Wu and Davis, 1993, Orr, 1997, Laurie, 1997, Price and Bouvier, 2002, Presgraves, 2002, Coyne and Orr, 2004); and the large X-effect, the observation that the X chromosome has a disproportionately large effect on hybrid sterility (Coyne and Orr, 1989, Coyne, 1992a, Presgraves, 2008). These patterns hold across a wide range of taxa, including female heterogametic (*ZW*) birds and Lepidoptera and male heterogametic (*XY*) plants, *Drosophila*, and mammals (Coyne and Orr, 1989, Coyne and Orr, 2004). We now know that these “two rules of speciation” (Coyne and Orr, 1989) are, in part, attributable to the rapid evolution of genetic factors that cause interspecific hybrid sterility on the X chromosome relative to the autosomes (Tao and Hartl, 2003, Moehring et al., 2007, Masly and Presgraves, 2007, Presgraves, 2008, Good et al., 2008). The relatively rapid accumulation of X-linked hybrid sterility factors is associated with reduced interspecific gene flow at X-linked *versus* autosomal loci {Muirhead, 2016 #3060}{Turissini, 2017 #3300}{Garrigan, 2012 #2715}. Overall, these patterns show that, for many taxa with heteromorphic sex chromosomes, the X chromosome plays a large and fundamental role in speciation.

Given the taxonomic breadth of Haldane’s rule, the large X-effect, and reduced interspecific gene flow on the X, understanding *why* the X chromosome accumulates hybrid incompatibilities faster than the rest of the genome is imperative. At least five explanations have been proposed: faster X evolution (Charlesworth et al., 1987), gene traffic (Moyle et al., 2010), disrupted sex chromosome regulation in the germline (Lifschytz and Lindsley, 1972), the evolutionary origination of incompatibilities in parapatry (Hollinger and Hermisson, 2017), and meiotic drive (Hurst and Pomiankowski, 1991, Frank, 1991).

Here we focus on the potential role of meiotic drive. The drive theory posits that sex chromosomes are more susceptible than autosomes to invasion by selfish meiotic drive (*sensu lato*) elements (Hurst and Pomiankowski, 1991, Frank, 1991). Sex-linked drive compromises fertility and distorts sex ratios, which leads to evolutionary arms races between drivers, unlinked suppressors, and linked enhancers (Lindholm et al., 2016, Presgraves, 2008, Meiklejohn and Tao, 2010). These arms races can contribute to the evolution of hybrid male sterility, in at least two ways. Normally-suppressed drive elements might be aberrantly expressed in the naïve genetic backgrounds of species hybrids, causing sterility rather than sex ratio distortion (Hurst and Pomiankowski, 1991, Frank, 1991). Alternatively, recurrent bouts of invasion, spread, and coevolution among drive, suppressor, and enhancer loci might cause interspecific divergence at these loci that secondarily cause hybrid sterility and map disproportionately to sex chromosomes (Presgraves, 2008, Meiklejohn and Tao, 2010).

Multiple lines of evidence support the plausibility of the drive theory. First, theoretical considerations and empirical evidence suggests that both active and suppressed sex chromosome meiotic drive systems are widespread in natural populations (Jaenike, 2001). Indeed, in one species, *Drosophila simulans*, at least three cryptic (normally suppressed) *sex-ratio* drive systems—Winters, Durham, and Paris—have been identified, involving distinct sets of X-linked drive loci and autosomal and/or Y-linked suppressors (Tao et al., 2001, Tao et al., 2007a, Tao et al., 2007b, Helleu et al., 2016). Second, loci involved in cryptic *sex-ratio* systems co-localize with hybrid male sterility loci in genetic mapping experiments (Tao et al., 2001, Zhang et al., 2015). Third, one of the two X-linked hybrid sterility genes identified to date also causes meiotic drive (Phadnis and Orr, 2008). These discoveries confirm that recurrent bouts of drive and suppression have occurred during the history of a single lineage and that cryptic drive genes can cause hybrid sterility. While these findings put the plausibility of the drive hypothesis beyond doubt, the question of its generality remains: what fraction of X-linked hybrid sterility factors evolved as a consequence of drive? Furthermore, the drive hypothesis (Hurst and Pomiankowski, 1991, Frank, 1991) assumes that populations evolve in strict allopatry (simple speciation) and/or that drive elements require population-specific genetic backgrounds for their activity. For populations that diverge with some level of gene flow (complex speciation), drive elements that are not population-specific can in principle migrate between species, thereby reducing divergence and undermining the evolution of hybrid sterility (Macaya-Sanz et al., 2011, Crespi and Nosil, 2013, Seehausen et al., 2014).

Here we investigate the special role of sex chromosomes in speciation with genetic mapping and population genomic analyses between *Drosophila mauritiana* and *D. simulans*. The human commensal species, *D. simulans*, originated on Madagascar, diverging from the sub-Saharan African species, *D. melanogaster*, ~3 Mya (Lachaise et al., 1988, Dean and Ballard, 2004, Baudry et al., 2006, Kopp, 2006, Ballard, 2004). The island-endemic species, *D. mauritiana*, originated on the Indian Ocean island of Mauritius, diverging from *D. simulans* ~240 Kya (Kliman et al., 2000, McDermott and Kliman, 2008, Garrigan et al., 2012). The two species are now isolated by geography— *D. simulans* has never been collected on Mauritius {David, 1989 #2349}— and by multiple incomplete reproductive incompatibilities, including asymmetric premating isolation (Coyne, 1992b), postmating-prezygotic isolation (Price, 1997), and intrinsic postzygotic isolation (F_1_ hybrid males are sterile, F_1_ hybrid females are fertile; (Lachaise et al., 1986)). Despite geographic and reproductive isolation, there is clear evidence for historical gene flow between the two species (Solignac and Monnerot, 1986, Solignac et al., 1986, Garrigan et al., 2012, Ballard, 2000a, Ballard, 2000b, Satta et al., 1988, Satta and Takahata, 1990). The X chromosome shows both an excess of factors causing hybrid male sterility (True et al., 1996b, Tao et al., 2003) and, correspondingly, a dearth of historical interspecific introgression (Garrigan et al., 2012). The rapdi accumulation of X-linked hybrid male sterility factors may have contributed to reduced X-linked gene flow, limiting exchangeability at sterility factors and genetically linked loci (Muirhead and Presgraves, 2016).

To begin to assess the role of drive in the evolution of X-linked hybrid male sterility between these two species, we performed genetic mapping experiments using genotype-by-sequencing of advanced-generation recombinant X-linked introgressions from *D. mauritiana* in an otherwise pure *D. simulans* genetic background. In parallel, we performed the first population genomics analyses of speciation between *D. mauritiana* and *D. simulans* to study the chromosomal distributions of interspecific divergence and introgression. These analyses lead to two discoveries regarding the role of meiotic drive in speciation. First, we find evidence for weak X-linked segregation distortion in hybrids, supporting the hypothesis that cryptic *sex-ratio* systems are common. Second, we report a surprising discovery concerning the role of drive during complex speciation: a now-cryptic sex ratio drive system recently introgressed between species and caused large selective sweeps in both species. As a result, this large X-linked region shows greatly reduced interspecific sequence divergence and an associated lack of hybrid male sterility factors. These findings suggest that the effects of selfish genetic elements on interspecific divergence and the accumulation of incompatibilities depends on the opportunity for drive systems to migrate between species during complex speciation.

## RESULTS

### Mapping X-linked hybrid male sterility

Multiple intervals on the X chromosome cause male sterility when introduced from *D. mauritiana* into *D. simulans* (True et al., 1996b, Maside et al., 1998). The number and identities of the causal factors, how they disrupt spermatogenesis, and the evolutionary forces that drove their interspecific divergence are unknown. We therefore generated a high-resolution genetic map of X-linked hybrid male sterility between the two species, with the ultimate aim of identifying a panel of sterility factors. We first introgressed eight X-linked *D. mauritiana* segments that together tile across ~85% of the euchromatic length of the X chromosome into a *D. simulans* genetic background (Figure 1A,B; Table 1). Each introgressed segment was marked by two co-dominant *P*[*w*^+^] insertions (True et al., 1996a) that serve as visible genetic markers. We introgressed these “2*P*” segments into the *D. simulans w*^XD1^ genetic background through >40 generations of repeated backcrossing (Figure 1A). Our ability to generate these introgression genotypes confirms that the distal 85% of the *D. mauritiana* X euchromatin carries no dominant factors that cause female sterility or lethality in a *D. simulans* genetic background (True et al., 1996b, Tao et al., 2003). All eight 2*P* introgression genotypes are however completely male-sterile, indicating that each of the introgressed regions contains one or more hybrid male sterility factors. Two pairs of introgression genotypes carry largely overlapping introgressed *D. mauritiana* segments and were combined for further analyses (2*P-*5a/b and 2*P*-6a/b, respectively; Figure 1B, Table 1).

**Figure 1.**
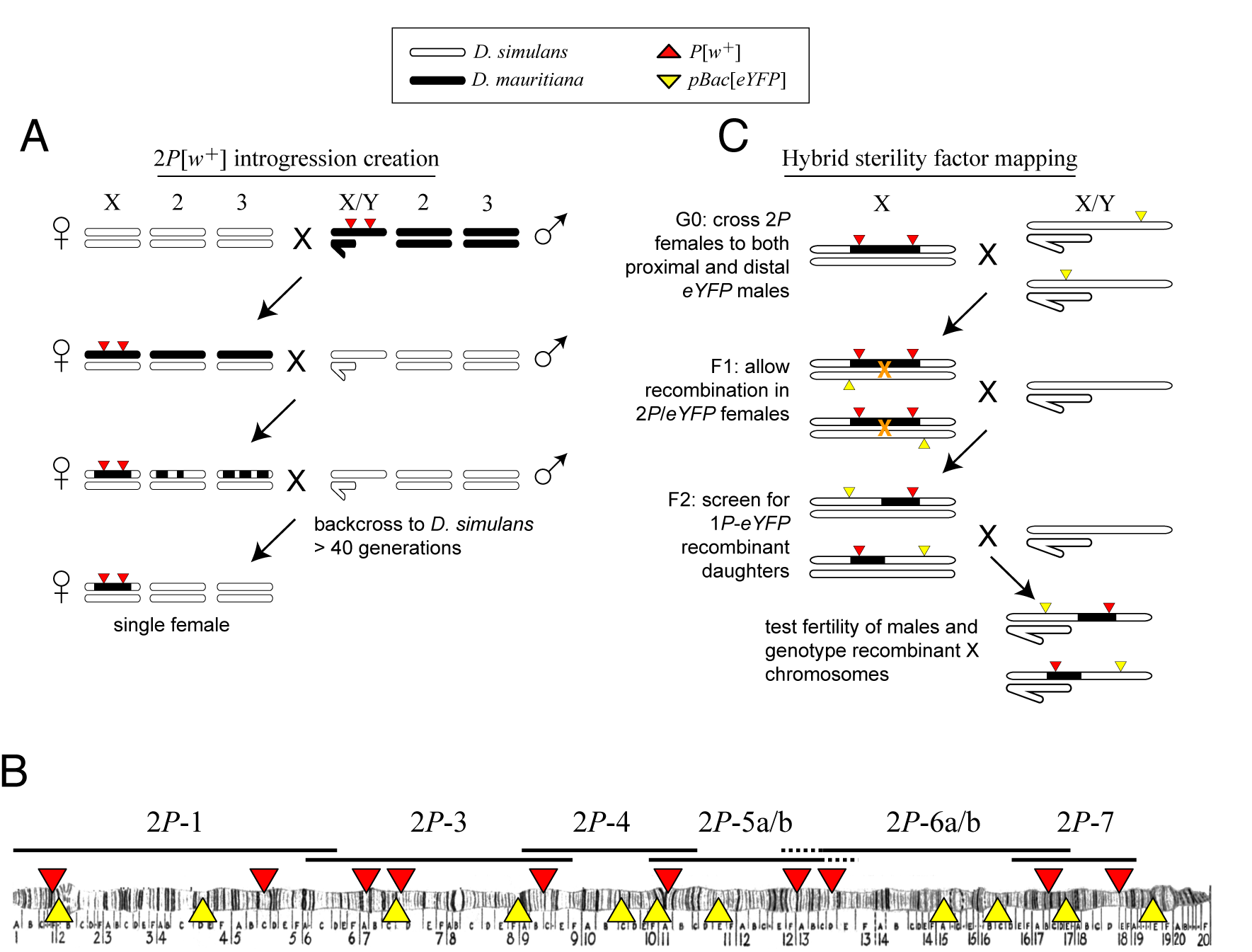
Crosses used to introgress eight regions of the *D. mauritiana* X chromosome into a *D. simulans* genome. (A) *D. mauritiana* “2*P*” lines were constructed by combining pairs of *P*-element insertions containing the miniwhite transgene (*P*[*w*^+^]; red triangles) distributed across the X chromosome. The *P*[*w*^+^] inserts are semi-dominant visible eye-color markers that permit discrimination of individuals carrying 0, 1 or 2 *P*[*w*^+^]. X-linked segments from *D. mauritiana* were introgressed into a *D. simulans* genetic background by backcrossing 2*P*[*w*^+^] hybrid females to *D. simulans w*^XD1^ males for over 40 generations. Each introgression line was then bottlenecked through a single female to eliminate segregating variation in the recombination breakpoints flanking the 2*P*[*w*^+^] interval. (B) Cytological map of the *D. melanogaster* X chromosome, indicating the locations of *P*[*w*^+^] and *pBac*[*eYFP*] transgene insertions. The extent of regions introgressed from *D. mauritiana* into *D. simulans* (*e.g.* 2*P*-1) are labeled above the map. Two pairs of introgression genotypes (2*P*-5a/b and 2*P*-6a/b) mostly overlap; the regions included in 2*P*-5b/2*P*-6b but not 2*P*-5a/2*P*-6a are indicated by dashed lines. (C) Meiotic mapping of sterility factors. 2*P*[*w*^+^] females were crossed to *D. simulans* strains carrying an X-linked *pBac*[*eYFP*] transgene (yellow triangles) that was used as an additional visible marker to score recombinant chromosomes. Recombinant X chromosomes with both *pBac*[*eYFP*] and a single *P*[*w*^+^] were chosen and assayed for male fertility. Recombinant chromosomes were generated using *pBac*[*eYFP*] markers both proximal and distal to each 2*P* introgression.

**Table 1.**
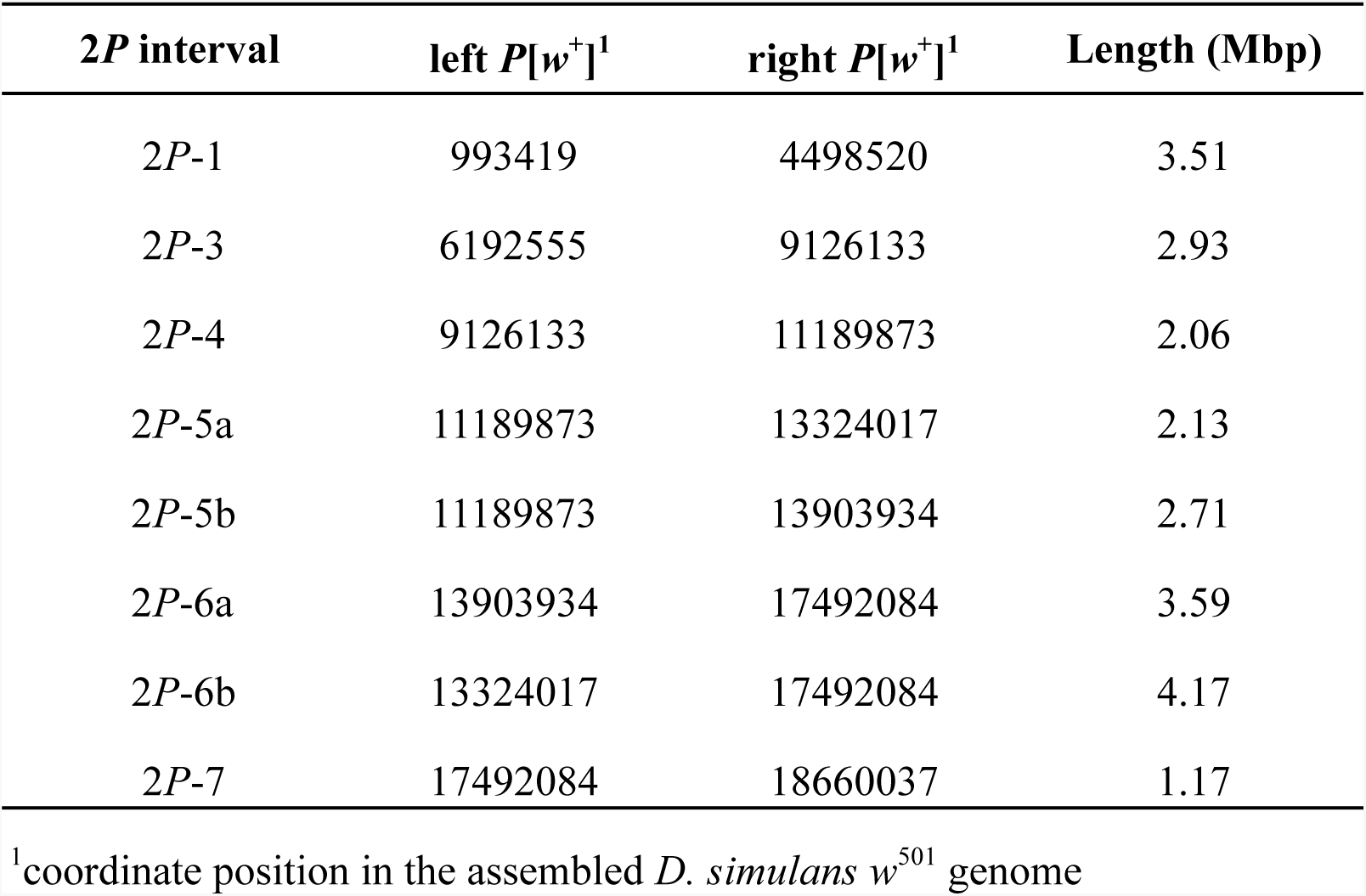
Locations and lengths of 2*P* intervals.

To determine the genetic basis of male sterility within each 2*P* interval, we generated recombinant introgressions using *D. simulans* strains carrying *pBac*[*eYFP*] visible markers (Stern et al., 2017) (Figure 1C). These crosses capture unique recombination events between *P*[*w*^+^] and *pBac*[*eYFP*] markers, allowing recombinant *D. mauritiana* introgressions (hereafter called 1*P-YFP*) to be propagated indefinitely through females without recombination via selection for the 1*P-YFP* genotype. From these 1*P-YFP* females, an unlimited number of replicate males carrying identical 1*P-YFP* recombinant introgressions can be generated, assayed for male fertility, and archived for genotyping (Figure 1C; see below). We assayed male fertility in at least ten individual males from each of 617 recombinant 1*P*-*YFP* genotypes (Table 2; see Methods). Across 1*P-YFP* genotypes, the mean number of offspring ranged from zero to 215 progeny; 238 (38.6%) were completely male-sterile, producing no offspring, and an additional 62 (10%) produced fewer than five offspring per male (Figure S1). Of the remaining 1*P-YFP* genotypes, 231 (37.4%) had intermediate fertility, and 86 (13.9%) had fertility indistinguishable from within-species controls (*P*_*t-test*_ > 0.01).

**Table 2.** Fertility and sex ratio pheuotypes for 1*P-YFP* recombinant genotypes.

| 2P interval | *n* tested | *n* sterile<sup>0</sup> | *n* fertile<sup>1</sup> | mean fertility<sup>2</sup> | % fertile<sup>1</sup> | mean SR<sup>2</sup> |
| --- | --- | --- | --- | --- | --- | --- |
| 2P-1 | 171 | 48 | 108 | 73.52 | 0.63 | 0.43 |
| 2P-3 | 97 | 12 | 64 | 67.38 | 0.66 | 0.45 |
| 2P-4 | 77 | 17 | 51 | 71.94 | 0.66 | 0.45 |
| 2P-5a/b | 92 | 23 | 53 | 68.15 | 0.58 | 0.51 |
| 2P-6a/b | 97 | 69 | 18 | 73.83 | 0.19 | 0.44 |
| 2P-7 | 83 | 69 | 9 | 133.00 | 0.11 | 0.47 |
| all 1P-YFP genotypes | 617 | 238 | 303 | 81.30 | 0.47 | 0.45 |

<sup>0</sup>genotypes where no male produced any offspring
<sup>1</sup>genotypes where at least one male produced at least five offspring
<sup>2</sup>among males producing more than four offspring

We determined high-resolution genotypes of 1*P*-*YFP* recombinant introgressions using multiplexed whole-genome sequencing (Andolfatto et al., 2011). After quality filtering, we obtained high-confidence genome-wide genotype information for 439 1*P*-*YFP* recombinant introgressions. No genotype showed evidence for any autosomal *D. mauritiana* alleles, confirming that the introgression scheme isolated X-linked *D. mauritiana* segments in a pure *D. simulans* autosomal genetic background (Figure S2). Recombinant 1*P*-*YFP* introgressions on the X chromosome ranged in size from 0.219 to 6.32 Mbp, with a mean length of 1.97 Mb (Table 3). Figure 2 shows the distribution of *D. mauritiana* introgression segments and their corresponding sterility phenotypes. Three large regions on the *D. mauritiana* X chromosome can be introgressed into *D. simulans* without severely negative effects on male fertility, indicating an absence of major hybrid male sterility factors in these regions (Figure 2). Conversely, we delineated four small regions (<700kb) that consistently and strongly reduced male fertility: 90% of replicate males with introgressions spanning these regions produce fewer than five offspring. Quantitative trait locus (QTL) analyses confirmed the existence of genetic variation among introgression genotypes that significantly affects male fertility (Figure 3). At least five QTL peaks are significant at *P* < 0.01 (permutation test). Most regions containing *D. mauritiana* alleles reduce the average number of progeny to <15. Two QTL peaks (2.5cM, and 29.3cM, Figure 3) appear to show higher fertility associated with the *D. mauritiana* allele than the *D. simulans* allele, but this is attributable to *D. mauritiana* sterility factors located at 12.6cM and ~35cM and the negative linkage disequilibrium that is generated across a 2*P* interval by our meiotic mapping approach (Figure 1C). Similar results are obtained with QTL analyses that treat each 2*P* interval as a separate mapping population (Figure S3).

**Table 3.** Distribution of 1*P-YFP* recombinant introgression lengths.

| 2P interval | sequenced | min size | mean size | max size |
| --- | --- | --- | --- | --- |
| 2P-1 | 129 | 295,225 | 2,617,833 | 6,322,871 |
| 2P-3 | 73 | 306,052 | 1,636,944 | 3,818,569 |
| 2P-4 | 55 | 226,018 | 1,482,659 | 2,917,578 |
| 2P-5 | 61 | 365,004 | 1,627,632 | 3,276,930 |
| 2P-6 | 55 | 692,350 | 2,400,499 | 4,764,204 |
| 2P-7 | 66 | 218,722 | 1,412,108 | 2,502,552 |
| total | 439 |  |  |  |

**Figure 2.**
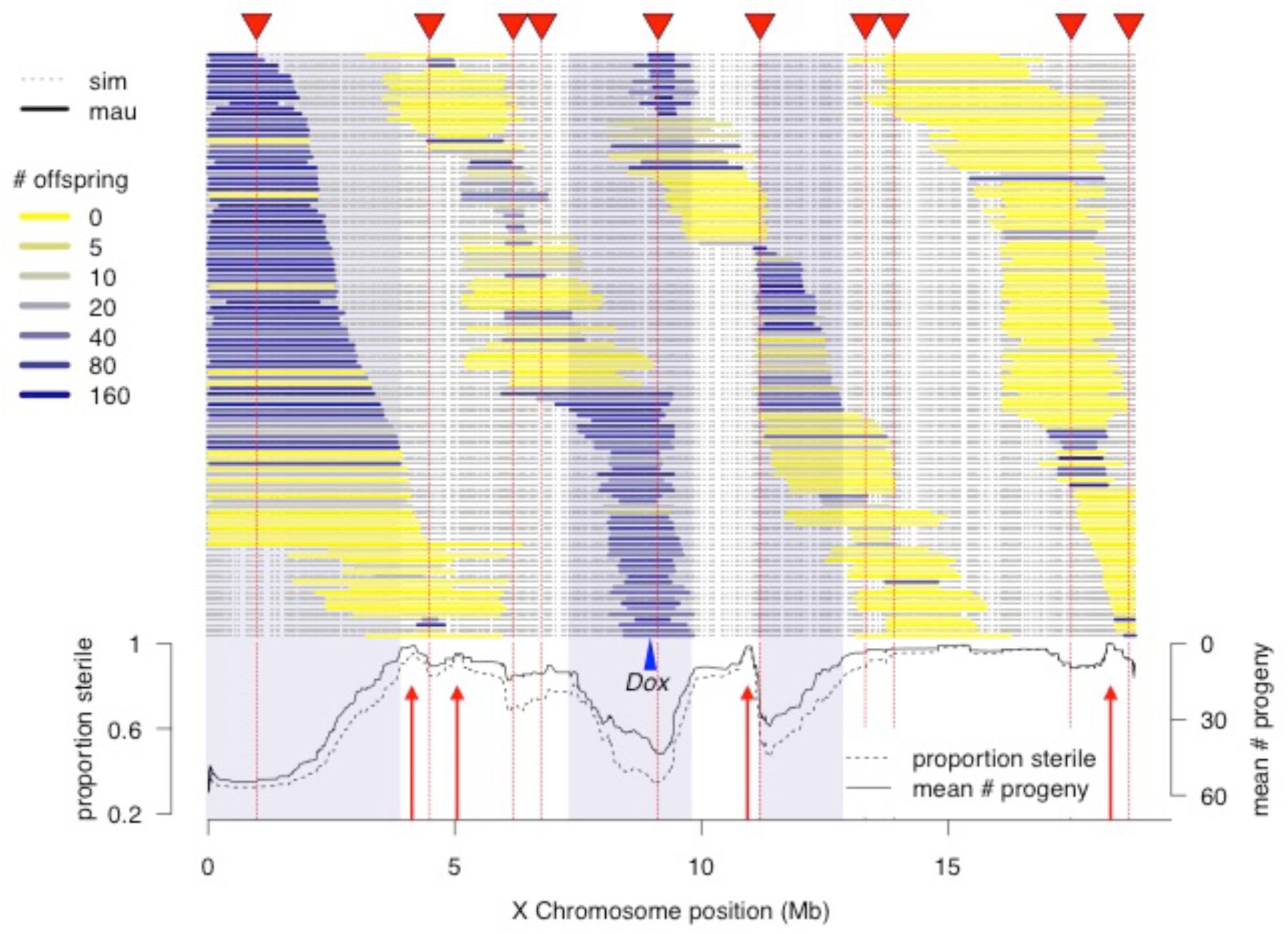
High-resolution genetic map of X-linked hybrid male sterility. Colored horizontal bars indicate the extent of introgressed *D. mauritiana* alleles for each recombinant 1*P-*YFP X chromosome. The color of each introgression indicates the mean fertility of ten replicate males carrying that 1*P-*YFP X chromosome. The three shaded areas indicate fertile regions within which *D. mauritiana* introgressions do not cause sterility, whereas the four red arrows indicate small candidate sterility regions. The blue arrowhead indicates the location of the *Dox/MDox* meiotic drive loci. Lines indicate the average number of offspring and average proportion of sterile males (defined as producing fewer than 5 offspring) for all 1*P-*YFP genotypes that carry *D. mauritiana* alleles at each genotyped SNP.

**Figure 3.**
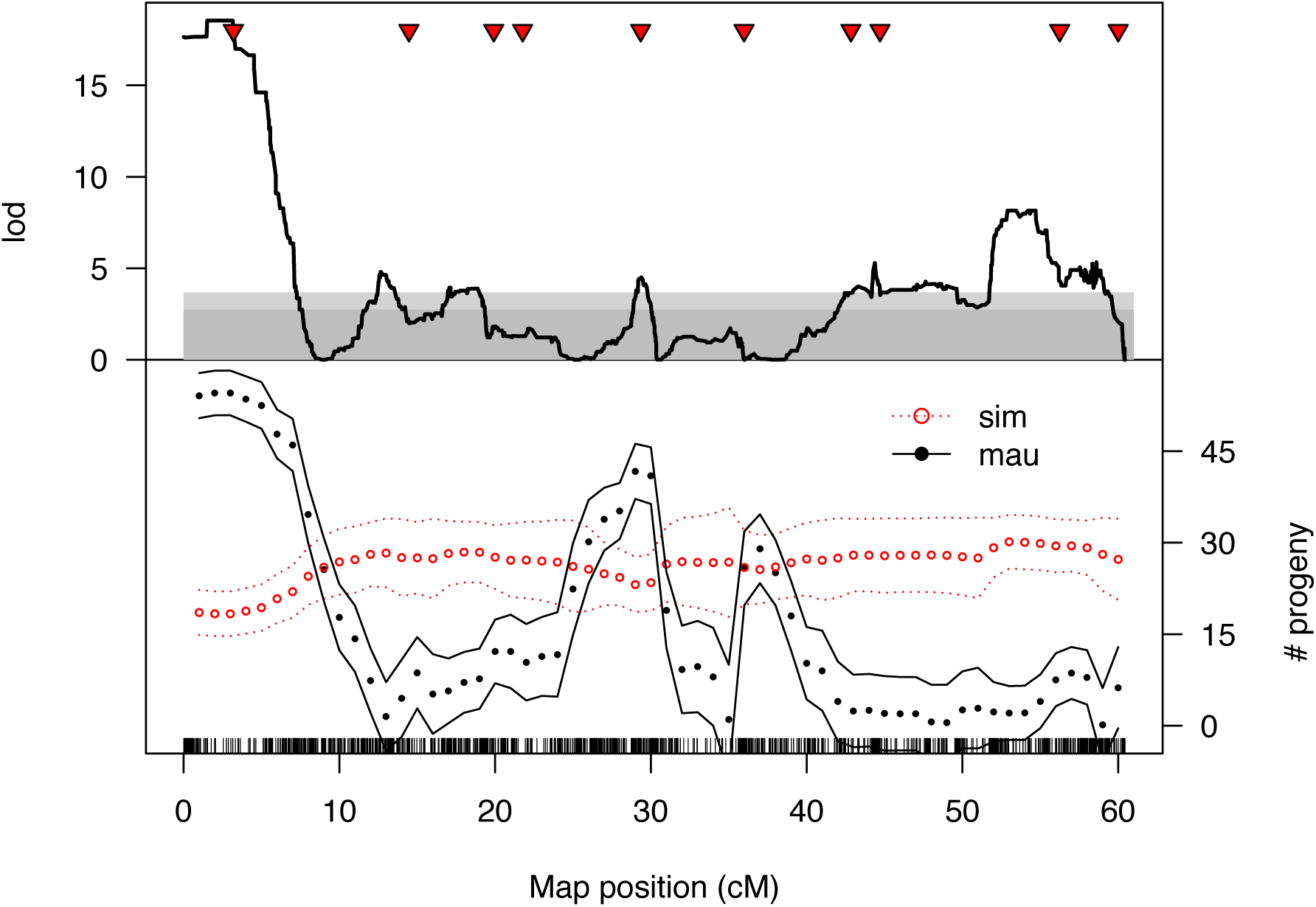
QTL analysis of male fertility. (A) Lod scores for a single-factor scan using all 1P-YFP introgression genotypes as a single mapping population. Dark and light gray regions indicate 5% and 1% significance thresholds, respectively, determined from 10.000 random permutations. (B) The estimated effects of *D. simulans* and *D. mauriticma* alleles at QTL placed every 1cM (bounding lines indicate 95% confidence intervals).

### Sex ratio distortion revealed through experimental introgression

Among fertile 1*P-YFP* males, progeny sex ratios were skewed toward a slight excess of sons: the mean proportion of daughters was 0.45. and 86% of fertile 1*P-YFP* genotypes (260/303) produced fewer than 50% daughters (Figure 4). However, these skewed sex ratios are at least partially attributable to effects of the *sim w*^XD1^ genetic background, as a similar male bias was observed among progeny of control *sim w*^XD1^ males (mean proportion females = 0.46. *n* = 35 sires, *t*-test *vs*. null hypothesis of 0.5. *P* = 0.005). Introgressed *D. mauritiana* alleles may modify this modest male bias — across all fertile introgression genotypes, there is a significant negative correlation between the length of the introgressed segment and the proportion of female progeny produced by that genotype (*r*=-0.31. *P*<0.0001). This effect seems to be independent of a relationship between the length of introgressed segments and fertility: while there is also a significant negative correlation between introgression length and mean number of progeny (*r*=-0.22. *P*<0.0001). there is not a significant correlation between mean number of progeny and progeny sex ratio (*P*=0.36). One interpretation of these results is that the *Y* chromosome of *sim w*^XD1^ causes weak segregation distortion, and the intensity of distortion is modified by X-linked alleles from *D. mauritiana*.

**Figure 4.**
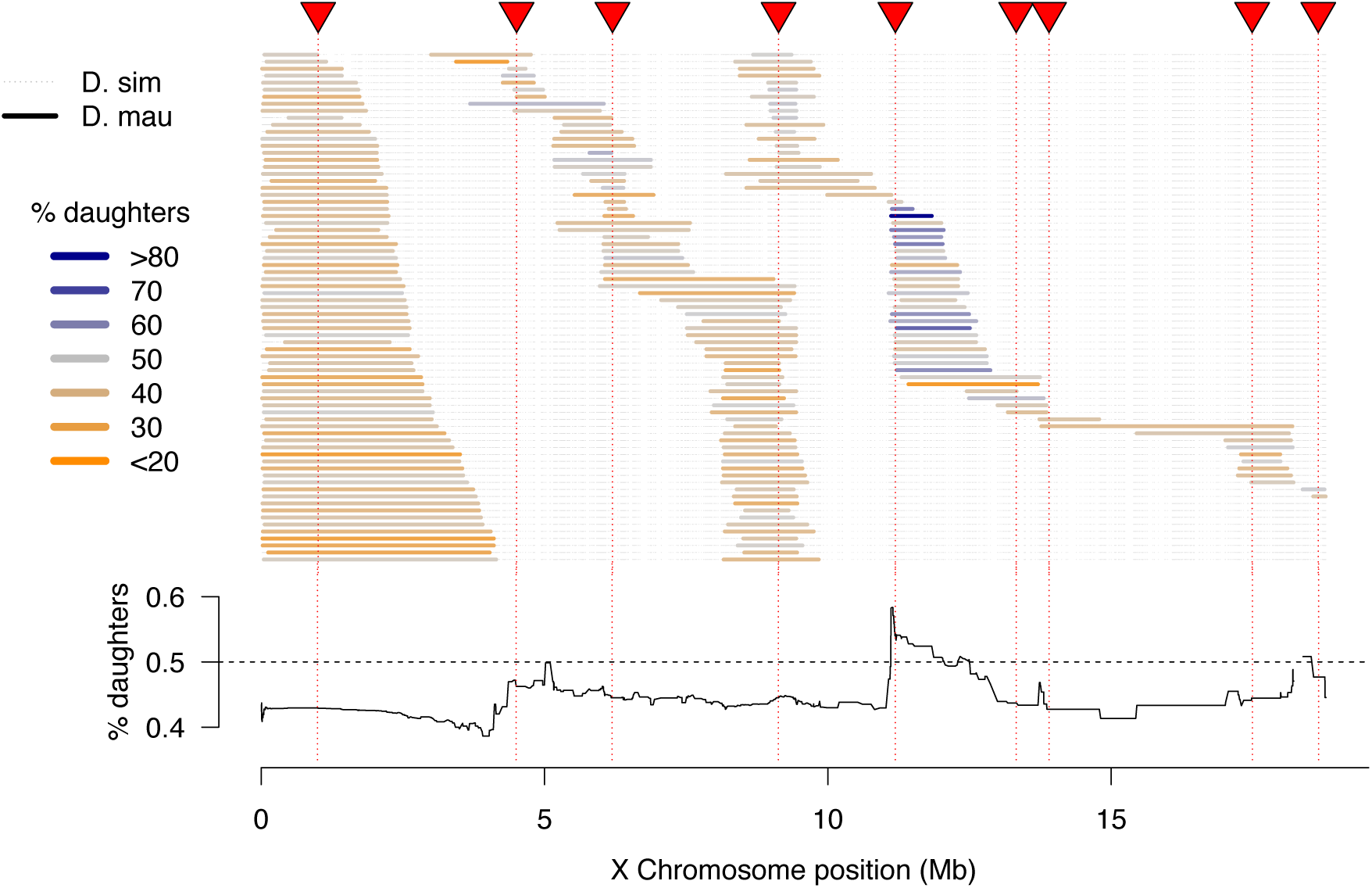
High-resolution map of progeny sex ratios among fertile 1*P-YFP* introgression male genotypes. Colored horizontal bars indicate the extent of introgressed *D. maurithma* alleles for each fertile recombinant 1*P-YFP* X chromosome. The color of each introgression indicates the sex-ratio of progeny from replicate males carrying that 1*P-YFP* X chromosome. The line below indicates the average progeny sex-ratio for all 1P-YFP genotypes that cany *D. mauritiana* alleles at each genotyped SNP.

Although the majority of fertile 1*P-YFP* genotypes sired male-biased progeny, introgressions that included the distal end of the 2*P*-5 region sired female-biased progeny (Figure 4). QTL analysis of progeny sex ratio confirms a significant peak in the distal portion of 2*P*-5 (Figure S.4). The estimated effect of this QTL on progeny sex ratios is 54.6% daughters for the *mauritiana* allele *versus* 42.5% daughters for the *simulans* allele. These results are consistent with the existence of a cryptic (normally-suppressed) X-linked drive allele in *D. mauritiana* that is released in a *D. simulans* genetic background-as the *D. mauritiana w*^12^ strain used to generate the IP introgressions produces slightly male-biased progeny sex-ratios using the same fertility assay (one male paired with three D. simulans *w*^XD1^ females, *n* = 10 sires, mean sex-ratio = 0.47. Mest vs. *D. simulans sim w*^XD1^ *P* = 0.4). This region of the X chromosome does not contain any previously mapped meiotic drive loci in *D. simulans* (Montchamp-Moreau et aL 2006. Tao et aL. 2007a. Helleu et aL. 2016). suggesting that these experiments have uncovered a novel cryptic drive locus and provide the first evidence of cryptic X-chromosome drive in *D. mauritiana*.

### Population genomics of speciation history

The high density of hybrid male sterility factors, and the presence of cryptic drive systems on the X chromosome is expected to influence patterns of gene flow between *D. mauritiana* and *D. simulans.* We therefore analyzed whole-genome variation within and between 10 *D. mauritiana* strains (Garrigan et al., 2014) and 21 *D. simulans* strains (Rogers et al., 2014), which allows us to characterize differentiation and identify genomic regions with histories of recent interspecific introgression. These analyses complement earlier studies that characterized interspecific divergence (Garrigan et al., 2012), polymorphism within *D. mauritiana* (Garrigan et al., 2014, Nolte et al., 2013), and polymorphism within *D. simulans* (Begun et al., 2007, Rogers et al., 2014). Below we present genome-wide population genetic analyses using non-overlapping 10-kb windows (see Methods).

### Polymorphism

Our genome-wide analyses provide multiple indicators that the island-endemic *D. mauritiana* has a smaller effective population size than *D. simulans* (Table 4), consistent with previous multi-locus analyses (Hey and Kliman, 1993, Kliman et al., 2000). Compared to *D. simulans*, total polymorphism (Nei and Li, 1979) in *D. mauritiana* is 32% lower on the X chromosome and 19% lower on the autosomes (Figure 5). The X/autosome ratio of polymorphism is thus lower in *D. mauritiana* (0.656) than in *D. simulans* (0.778) and lower than the ¾ expected for a random mating population with a 1:1 sex ratio (Garrigan et al., 2014). A substantial fraction of extant polymorphisms in both species arose in their common ancestor, reflecting the relatively recent speciation event and the fact that both species have large effective population sizes (see Methods). Compared to *D. simulans*, however, *D. mauritiana* has retained 74.4% as many ancestral polymorphisms and accumulated just 46.3% as many derived polymorphisms. The site frequency spectra (Tajima, 1989) in *D. mauritiana* are less skewed towards rare variants than in *D. simulans*, and average linkage disequilibrium (Kelly, 1997) is >2-fold higher. Overall, these findings show that, relative to *D. simulans*, *D. mauritiana* has lower nucleotide diversity; retained fewer ancestral SNPs; accumulated fewer derived SNPs; a less negatively skewed site frequency spectrum; and greater linkage disequilibrium— all patterns consistent with a historically smaller effective population size in *D. mauritiana* than in *D. simulans*.

**Table 4.**
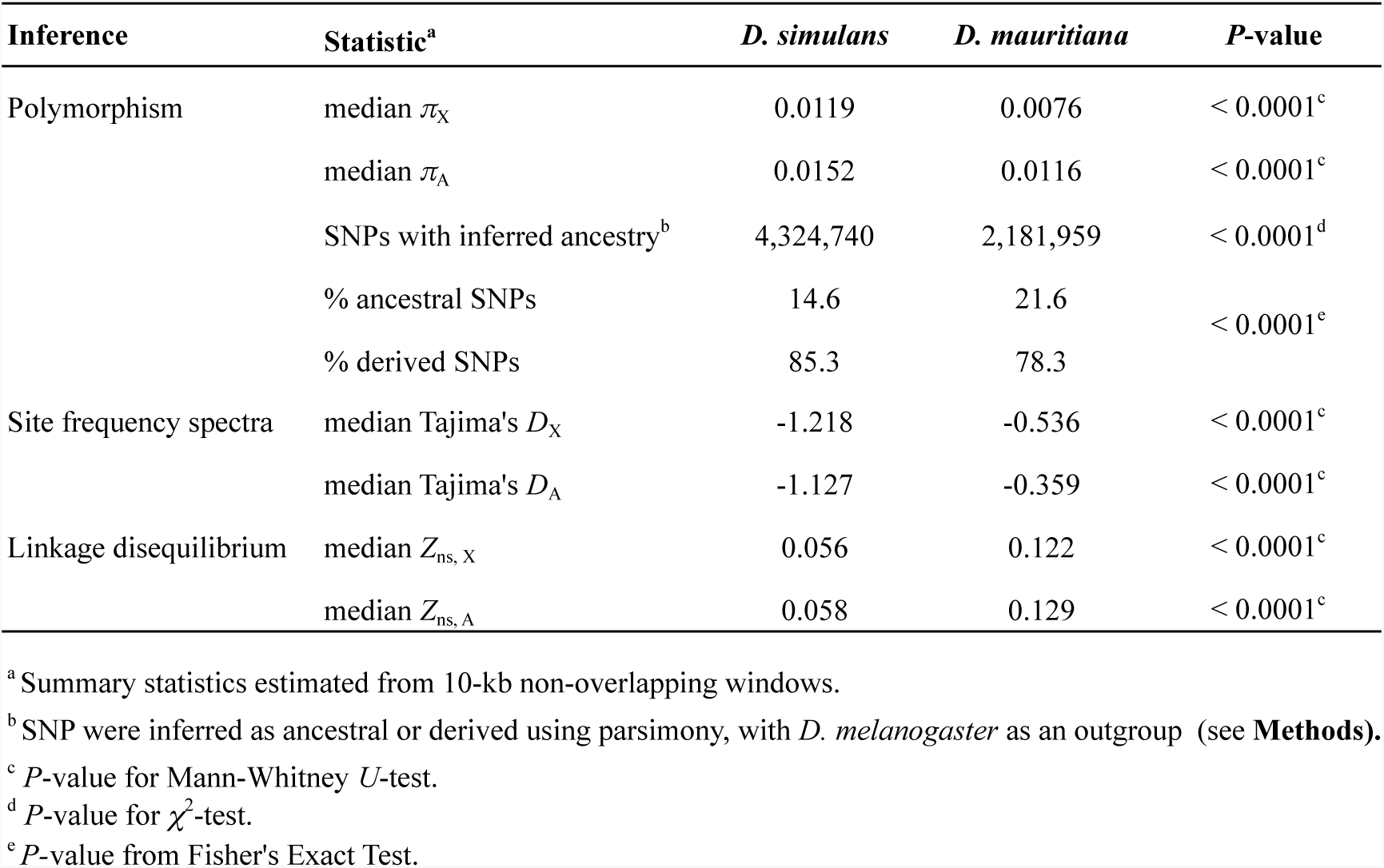
Population genomics summary statistics.

**Figure 5.**
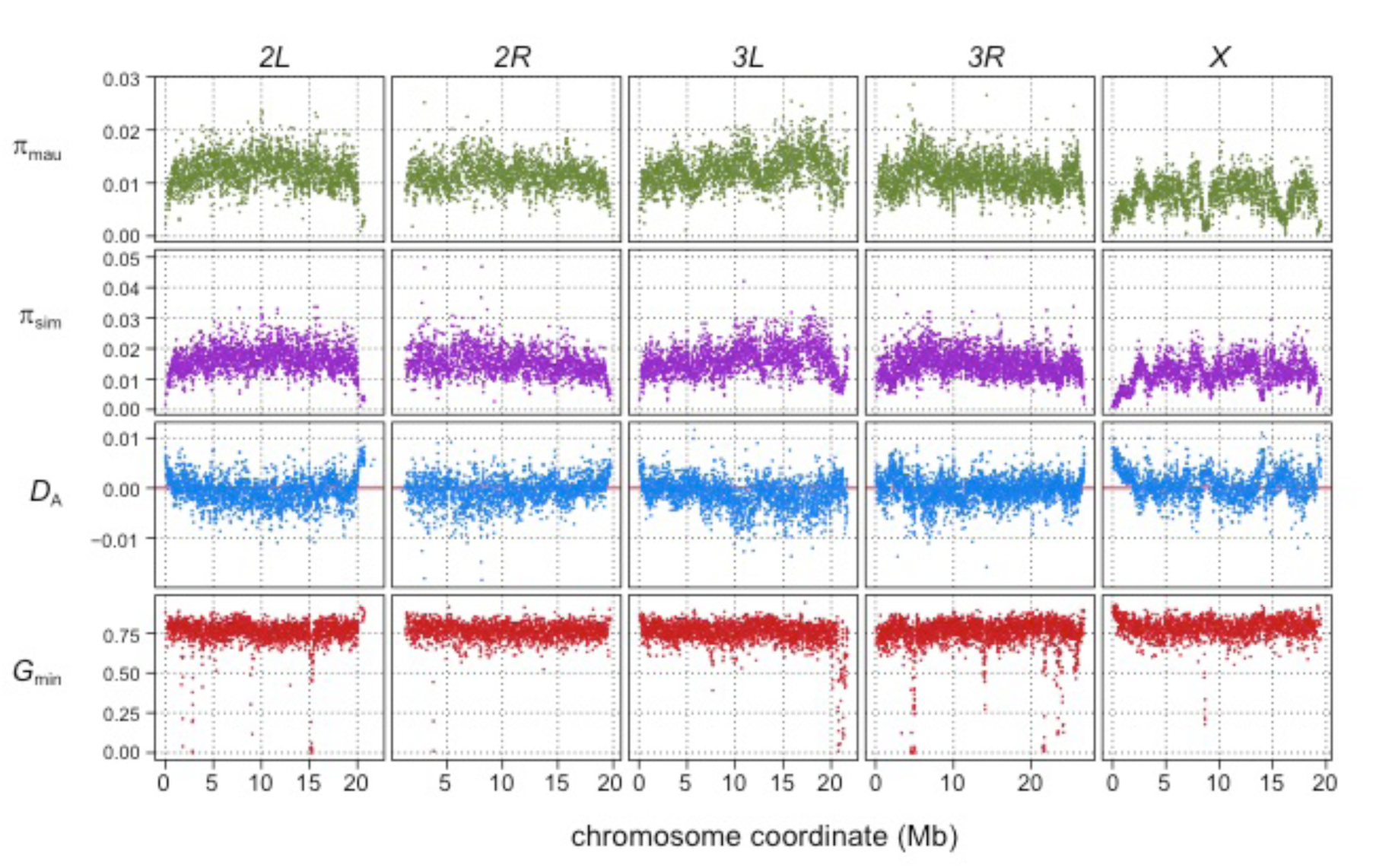
Population genomic scans for polymorphism, divergence, and introgression in 10-kb windows. The rows of panels show: nucleotide diversity for a sample of 10 inbred strains of *D. mauritiana* (π_mau_, green dots) and 20 inbred strains of *D. simulans* (π_sim_, purple dots); nucleotide divergence scaled by within-species polymorphism (blue dots); and *G*_min_ (red dots), the ratio of the minimum number of nucleotide differences per site between *D. mauritiana* and *D. simulans* to the average number of differences per site, a summary statistic that is sensitive to introgression. Panels correspond to each major chromosome arm, with genome coordinates on the *x*-axis.

### Divergence and differentiation

Net divergence levels between species are comparable to diversity levels within species. The median number of pairwise differences per site (*D*_*XY*_) between the two species, estimated in non-overlapping 10-kb windows, is 0.010 for the X chromosome and 0.013 for the autosomes. However, as the X chromosome has lower levels of polymorphism within species, the median net divergence (*D*_*A*_) between species is 0.0007 for the X and −0.0005 for the autosomes (a negative value of *D*_*A*_ on the autosomes occurs because, on average, levels of within-species polymorphism exceed levels of between-species divergence). Allele frequency differentiation is also higher for the X chromosome (median *F*_ST_=0.378) than the autosomes (median *F*_ST_=0.279, *P*_*MWU*_<2.2e-16). These estimates imply that, for X-linked and autosomal loci, the mean within-species sample coalescence times are 1.65- and 2.58-fold deeper than the species divergence time, respectively (Slatkin, 1993). The genetic basis of hybrid incompatibilities is therefore nested within the much deeper collective genealogical history of most genetic variation in *D. simulans* and *D. mauritiana*.

### Recent interspecific gene flow and introgression

Gene flow between *D. mauritiana* and *D. simulans* has been rare during their speciation history, with an apparent recent increase (Garrigan et al., 2012). To identify genomic regions that have introgressed between species in the very recent past, we used the *G*_min_ statistic— the ratio of the minimum pairwise sequence distance between species to the average pairwise distance between species (min[*D*_*XY*_]/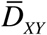; (Geneva et al., 2015). As populations diverge without gene flow, all loci in the genome gradually approach reciprocal monophyly, leaving just one ancestral lineage from each population available for coalescence in the ancestral population. Consequently, the minimum distance (numerator) equals the mean pairwise distance (denominator), causing *G*_min_→1 with zero variance. Conversely, *G*_min_ is small when the minimum distance is small relative to the mean pairwise distance. *G*_min_ is therefore sensitive to genealogical distortions resulting from recent gene flow, particularly when the introgressed sequence is at low to intermediate population frequency (Geneva et al., 2015). Between *D. mauritiana* and *D. simulans*, median *G*_min_ (± median absolute deviation) estimated for 10-kb windows across the major chromosome arms ranges from 0.761 ± 0.0537 for *3L* to 0.785 ± 0.0531 for the *X* (Figure 5; Kruskal-Wallis test, *P*<2.2e-16).

To identify 10-kb outlier windows that have genealogical histories inconsistent with strict allopatric divergence, we used a Monte Carlo simulation procedure that assumes a constant species divergence time across all 10-kb intervals, separately for the X and the autosomes (see Methods). In total, 196 of the 10,443 10-kb windows (1.9%) have a more recent common ancestry between *D. mauritiana* and *D. simulans* than expected under a strict allopatric divergence model, as indicated by significantly low values of *G*_min_ (*P* ≤0.001). As *G*_min_ is a ratio, significantly small *G*_min_ values could result from unusually small numerators (minimum *D*_*XY*_) or unusually large denominators (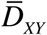). We find that 10-kb windows with significant *G*_min_ values have smaller median minimum *D*_*XY*_ (0.0056 in introgression windows *versus* 0.0094 genome-wide, *P*_MWU_ < 0.0001) as well as *smaller* median 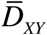 (0.0110 in introgression windows *versus* 0.0124 genome-wide *P*_MWU_ < 0.0001), indicating that the significant *G*_min_ values are due to unusually small minimum *D*_*XY*_ values. Introgression windows are 4.4-fold underrepresented on the X chromosome: only nine of 1833 10-kb windows on the X chromosome (0.49%) have significant *G*_min_ values *versus* 187 of 8414 10-kb windows on the autosomes (2.17%; Fisher’s exact test *P*=6.5e-08). However, not all 10-kb introgression windows are independent: 169 of the 196 significant 10-kb windows (86.2%) can be arrayed into contiguous (or nearly contiguous) genomic regions (see Methods). As a result, we infer 27 small (10-kb) introgressions and 21 larger introgressions ranging in size from 20 kb to 280 kb (**Table S2**). Of these 48 total introgressions, only one is on the X chromosome and 47 are on autosomes (*X*^2^-test, *P*=0.0124). The lengths of these introgressed haplotypes have been eroded by recombination over time in the receiving population: longer, presumably younger, introgressions have smaller minimum *D*_xy_ values (Spearman *ρ*=-0.526, *P*=0.00012). However, variation in local recombination rate has not been an important factor affecting introgression lengths (Spearman *ρ*=-0.0412, *P*=0.781).

### Interspecific introgression of the cryptic Winters *sex ratio* drive system

The single introgression detected on the X chromosome corresponds to a 130-kb region that comprises eight protein-coding genes plus the Winters *sex ratio* meiotic drive genes, *Distorter on the X* (*Dox*) and, its progenitor gene, *Mother of Dox* (*MDox*) (Tao et al., 2007a) (Figure 6). The median *G*_min_ value across this 130-kb region is 0.333, a ~2.4-fold reduction relative to background *G*_min_ on the X chromosome (*P*_MWU_*<*0.0001). In *D. simulans*, when unsuppressed, *MDox* and *Dox* cause biased transmission of the X chromosome, with male carriers siring excess daughters (Tao et al., 2007b). These drivers are suppressed by an autosomal gene, *Not much yin* (*Nmy*), which is a retrotransposed copy of *Dox*, via a putative RNA-interference mechanism (Tao et al., 2007b). In non-African *D. simulans* populations, *Dox, MDox*, and *Nmy* are nearly fixed, although haplotypes lacking the genes segregate at low frequencies (Kingan et al., 2010). All three loci have histories consistent with selective sweeps due to the presumed transmission advantage at *MDox* and *Dox* and the associated selective advantage of suppressing drive and restoring equal sex ratios at *Nmy* (Kingan et al., 2010).

**Figure 6.**
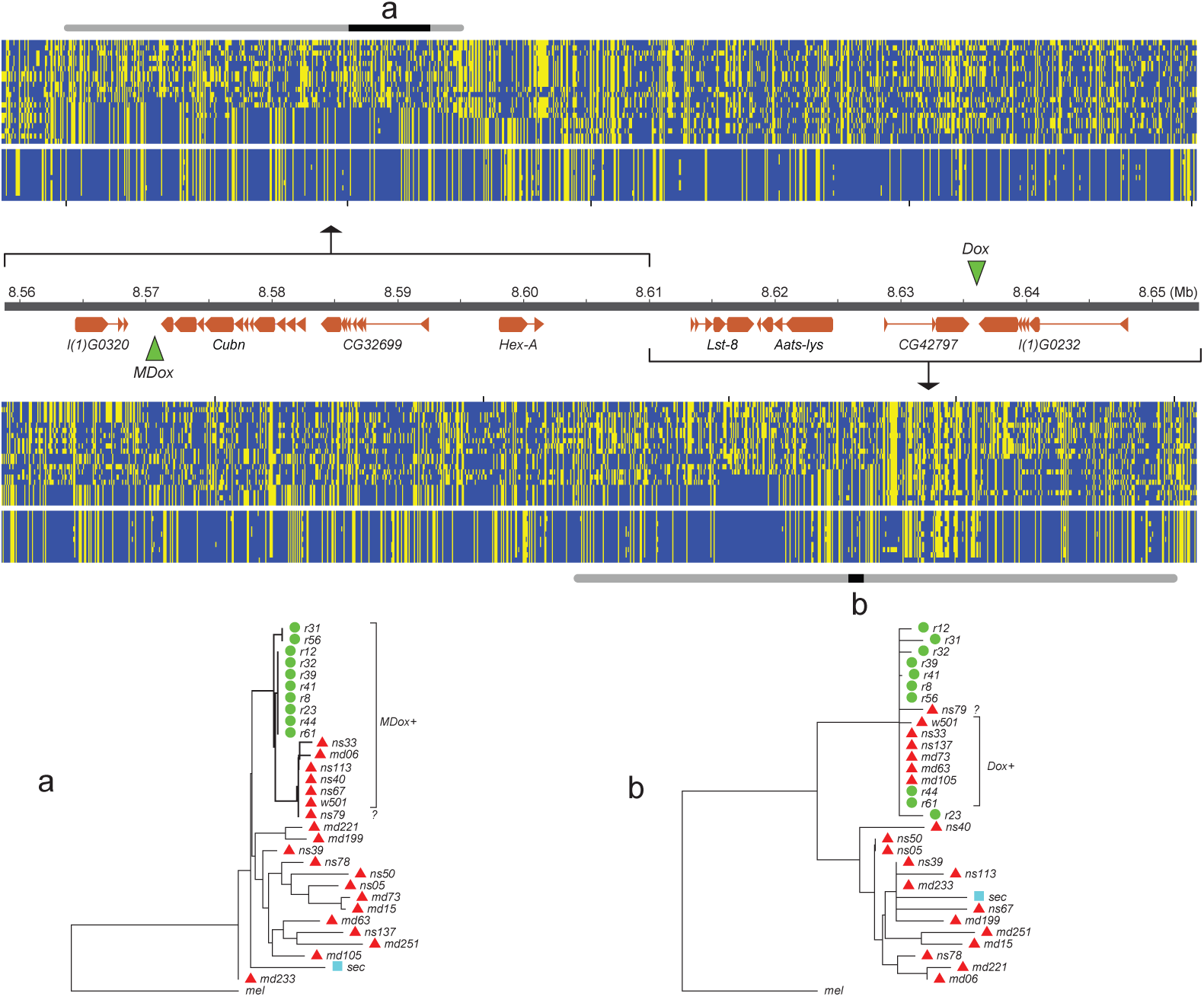
Natural introgression of the *MDox-Dox* region of the X chromosome. The top half of the figure shows two DNA polymorphism tables: the top table corresponds to the *MDox* region, and the bottom corresponds to the *Dox* region. Within the tables, yellow squares denote the derived nucleotide state, and blue squares indicate the ancestral state. The top 20 rows of each table correspond to the *D. simulans* samples, and the bottom 10 rows correspond to the *D. mauritiana* samples. The genome map between the polymorphism tables shows gene models for the region (orange boxes) and the locations of the *MDox* and *Dox* genes (green triangles). The grey bars on the top and bottom of the polymorphism tables mark the sites that occur within an introgressed region, with black sections labeled “a” and “b”, which mark the location of the *MDox* and *Dox* genes, respectively. The bottom panels of the figure show two maximum likelihood phylogenetic trees labeled “a” and “b”, corresponding to the *MDox* and *Dox* regions.

Maximum-likelihood phylogenetic trees for the 130-kb *MDox-Dox* region show reduced diversity within *D. mauritiana* and reduced divergence between species (Figure 6). Among the 10 *D. mauritiana* sequences, nucleotide diversity is just 24% (*π*=0.0018) of background diversity levels on the X chromosome, corresponding to a massive selective sweep in the *D. mauritiana* genome (*P*_MWU_<0.0001; see also (Nolte et al., 2013, Garrigan et al., 2014)). The distribution of variability among haplotypes in the *D. simulans* samples is consistent with a parallel, albeit incomplete, selective sweep (Figure 6). The most extreme 10-kb window within the 130-kb region has a minimum *D_XY_* value (=0.00087) that is 92% smaller than the X chromosome-wide 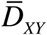, implying that introgression occurred in the very recent past.

To determine if the *MDox* and/or *Dox* drive elements are associated with introgression between species and the selective sweeps within each species, we determined *MDox* and *Dox* presence/absence status for each line using diagnostic restriction digests (see Methods). Previous work shows that *MDox* and *Dox* are nearly fixed among *D. simulans* samples collected outside of Africa (Kingan et al., 2010). However, among our 19 African samples (9 Madagascar, 10 Kenya), we find that the drivers are at lower frequency: five have *MDox* (26%), five have *Dox* (26%), and only one has both genes (5%; NS33; **Table S2**). Despite these low frequencies, *MDox* and *Dox* are overrepresented among the haplotypes shared between species: 6 of the 7 shared haplotypes have *MDox* and/or *Dox* (Fisher’s Exact *P*_FET_=0.0018), and 2 of the 7 possess both drivers (*P*_FET_=0.0158; *n*=19 African samples, plus the reference strain, *D. simulans w*^501^, which has both). In *D. mauritiana*, all 10 lines have *MDox*, but only two have *Dox* (Figure 6; **Table S6**). RT-PCR confirms that *MDox* is expressed in testes from both species (see Methods). These findings provide support for the notion that *Dox* and (transcriptionally active) *MDox* genes mediated the introgression and parallel sweeps.

The large *MDox-Dox* introgression, and its associated sweep co-localize with one of the three regions of the X chromosome that, in our mapping experiments, fails to cause sterility when introgressed from *D. mauritiana* into *D. simulans* (Figure 2). These observations suggest that a driving haplotype moved between species and swept to high frequency in *D. simulans* or fixation in *D. mauritiana*, thereby reducing local sequence divergence. This discovery has two implications. First, the *MDox-Dox* region is the only locus on the X chromosome to have recently escaped from its linked hybrid incompatibility factors and introgressed between species. Second, by sweeping to high frequency or fixation, the *MDox-Dox* drive element region reduced local divergence between species and, incidentally, undermined the accumulation of genetic incompatibilities that might cause hybrid male sterility.

## DISCUSSION

Our combined genetic and population genomics analysis of speciation between *D. mauritiana* and *D. simulans* yields three findings. First, we confirm the rapid accumulation of X-linked hybrid male sterility between these species and map four major sterility factors to small (<700 kb) intervals (Figure 2). Second, we find that very recent natural introgression has occurred between these species, albeit almost exclusively on the autosomes, consistent with a large X-effect on gene flow (**Table S.1**). Third, we discover new roles for meiotic drive during the history of speciation between these species. Some drive seems to have contributed to functional divergence between species: one region of the *D. mauritiana* X chromosome appears to cause segregation distortion in a *D. simulans* genetic background. In contrast, the well-characterized X-linked Winters *sex ratio* distorters, *MDox* and *Dox*, have clearly migrated between species, reducing local interspecific divergence. Together, these findings suggest that genetic conflict may both promote as well as undermine the special role of sex chromosomes in speciation.

### Genetic basis of X-linked hybrid male sterility

Our genetic analyses were initiated by introgression of six different regions of the *D. mauritiana X* chromosome into a pure *D. simulans* genetic background. All six regions cause complete hybrid male sterility and therefore carry at least one, or a combination of, *D. mauritiana* allele(s) that disrupts spermatogenesis due to incompatibilities with X-linked, Y-linked, or autosomal *D. simulans* alleles. Only three large (>2Mb) regions of the *D. mauritiana X* are readily exchangeable between species, permitting male fertility in a *D. simulans* genome. Thus, after only ~250,000 years, sufficient X-linked hybrid male sterility has accumulated to render most of the *D. mauritiana X* chromosome male-sterile on a *D. simulans* genetic background (True et al., 1996b). Most of the *D. mauritiana X* chromosome is male-sterile in a *D. sechellia* genome as well (Masly and Presgraves, 2007).

We were able to define four small regions (<700kb), each sufficient to cause complete male sterility (Figure 2), suggesting that these may contain single, strong sterility factors. We also find a large region spanning most of 2*P*-6 from which we were unable to recover fertile 1*P*-*YFP* recombinants. We infer that 2*P*-6 contains a minimum of two strong sterility regions, one tightly linked to each of the flanking *P-*elements (Figure 3). While our 2*P* mapping scheme is designed to facilitate the identification of male sterility factors, the 2*P*-6 interval highlights one of its limitations: in regions like 2*P*-6, for which strong sterility factors are very close to both flanking *P*-elements, we cannot determine how many additional sterility factors might localize to the middle of the interval. The present experiments therefore provide only a minimum estimate of the total hybrid male sterility factors on the X chromosome. We tentatively conclude that, within the fraction of the *D. mauritiana X* chromosome investigated, there are at least six genetically separable regions, each individually sufficient to cause virtually complete male sterility. It is worth noting these experimental approaches detect relatively large-effect sterility factors under a single set of laboratory conditions. There are likely many hybrid male sterility factors of smaller effect, generally neglected in the lab but easily detected by selection in natural populations and thus able to affect the probability of migration at linked loci.

### Genomic signatures of complex speciation with gene flow

The two species studied here are allopatric: *D. simulans* has never been reported on Mauritius, and *D. mauritiana* has never been found anywhere other than Mauritius (David et al., 1989, Legrand et al., 2011). *D. mauritiana* appears to have originated from a *D. simulans*-like ancestor, probably from Madagascar, that migrated and established a population on Mauritius (Hey and Kliman, 1993, Kliman et al., 2000). Our characterization of genome-wide variation within and between *D. mauritiana* and *D. simulans* confirms a coalescent history that reaches considerably deeper into the past than the inferred species split time of ~250,000 years (Hey and Kliman, 1993, Kliman et al., 2000). Nested within this largely shared coalescent history, many functional differences have evolved between the two species, including extreme ones that mediate large-effect hybrid incompatibilities. The signatures of gene flow found in the genomes of these species imply recurrent bouts of migration and interbreeding. To introgress between species, immigrating foreign haplotypes must escape their locally disfavored chromosomal backgrounds by recombination before being eliminated by selection against linked incompatibilities and locally maladaptive alleles (Petry, 1983, Bengtsson, 1985, Barton and Bengtsson, 1986). Conditional on escape, the lengths of foreign haplotypes will be subject to gradual erosion by recombination with the resident genetic background.

Here, and in previous work (Garrigan et al., 2012), we detect evidence consistent with weak migration: 2-5% of the genome shows evidence of introgression between *D. simulans* and *D. mauritiana* during their recent history. Our population genomic analysis of polymorphism and divergence using the *G*_min_ statistic identified 48 segregating foreign haplotypes. We find evidence that the genomic locations and lengths of introgressed foreign haplotypes have been shaped by selection and recombination in the receiving population, respectively. First, selection has likely affected the genomic distribution of foreign haplotypes: only one of the 48 introgressions occurs on the X chromosome. The opportunity for foreign haplotypes to escape the X chromosome via recombination is more constrained than on the autosomes, as the X has a higher density of incompatible alleles, and hemizygous selection eliminates foreign X-linked haplotypes more quickly (Muirhead and Presgraves, 2016). Second, after escaping locally deleterious chromosomal backgrounds, recombination has eroded the lengths of foreign haplotypes over time: recently introgressed, and hence less diverged, haplotypes tend to be longer. It is worth noting here that the 10-kb windows used for our *G*_min_ scan for foreign haplotypes almost certainly fails to identify smaller and/or older introgressions.

### Meiotic drive and complex speciation

The drive theory posits that hybrid incompatibilities accumulate as incidental by-products of recurrent bouts of meiotic drive and suppression (Hurst and Pomiankowski, 1991, Frank, 1991). Our mapping experiments provide no direct evidence in support of this theory in *D. mauritiana* and *D.simulans*, as no hybrid male sterility loci co-localized with sex-ratio loci. Our genetic mapping experiments have however provided further evidence for the accumulation of cryptic *sex-ratio* drive systems. We mapped a small region of the *D. mauritiana X* that, when introgressed into a naïve *D. simulans* genetic background, causes weak segregation distortion resulting in female-biased progeny sex ratios (Figure 4). As the *D. mauritiana* X-drive locus does not map to the location of any of the three cryptic drivers known from *D. simulans*, we infer that it may be a new, previously undiscovered drive system in *D. mauritiana*.

Across *D. simulans* and *D. mauritiana*, four cryptic drive systems have been identified so far: two X-drive systems in *D. simulans* (Paris and Durham); one X-drive system in *D. mauritiana* (see above); and one X-drive system found in both species (Winters; see below). We regard this as a minimum for several reasons. First, weak segregation distortion that may be significant in natural populations can go undetected in laboratory experiments. Second, cryptic drive systems may not be fixed within species, and we have only surveyed genotypes derived from one strain each of *D. mauritiana* and *D. simulans*. Third, no study has yet comprehensively assayed *D. simulans* material introgressed into a *D. mauritiana* genetic background. Finally, some cryptic drive systems might go to fixation and then simply die because once fixed (or suppressed), a driver is in a race: either suffer mutational decay or acquire a mutation that confers a new bout of drive. These considerations— and the discovery of multiple alternative cryptic drive systems in closely related species— imply that sex chromosome drive is not infrequent during the history of species divergence (Jaenike, 2001).

We have found that the Winters sex-ratio drivers, *MDox* and *Dox*, have moved between these two species. This discovery highlights an implicit assumption of the drive theory of the large X-effect—namely that species evolve in strict allopatry. With gene flow, drive elements (and other selfish genes) have the opportunity to jump species boundaries and undermine divergence in a process analogous to adaptive introgression (Seehausen et al., 2014, Crespi and Nosil, 2013). The *t*-haplotype has, for instance, introgressed between sub-species of house mouse, *Mus musculus* (Macaya-Sanz et al., 2011). Between *D. mauritiana* and *D. simulans*, the *G*_min_ statistic and the genealogies associated with the *MDox-Dox* introgressed haplotype (Figure 6) are agnostic on the direction of introgression. Nonetheless, the finding that a drive element crossed a species boundary has important implications for the drive theory explanation of Haldane’s rule and the large X-effect. For *MDox* and *Dox* to introgress between species, three things must be true: (1) neither *MDox* nor *Dox* alleles from the donor species caused male sterility in the recipient species; (2) no X-linked hybrid male sterility factors are so tightly linked to *MDox* and *Dox* as to prevent their eventual escape by recombination into the recipient species genetic background; and (3) any sterility factors located within the introgressed region of the recipient X will have been replaced by foreign alleles. Together, these inferences suggest that a selfish drive system was able to invade a new species by *not* causing male sterility and, for one X-linked region, may have impeded or undone the evolution of hybrid male sterility.

## METHODS

### Drosophila husbandry and genetics

All *Drosophila* crosses and phenotyping were done in parallel in two locations, using standard cornmeal media (Rochester, NY) or minimal cornmeal media (Bloomington, IN) at room temperature (23-25C). We constructed *D. mauritiana* “2*P*” lines that carry pairs of X-linked *P*-element insertions that contain the mini-*white* transgene (*P*[*w*^+^]) (True et al., 1996a) which serve as semi-dominant visible genetic eye-color markers and allow us to distinguish individuals carrying 0, 1 or 2 *P*[*w*^+^]. These “2*P*” regions were then introgressed into the *D. simulans w*^XD1^ genetic background through more than 40 generations of repeated backcrossing while following the two *P*[*w*^+^] insertions (Figure 1A). Each 2*P* introgression line was then bottlenecked through a single female to eliminate segregating variation in the recombination breakpoints flanking the 2*P*[*w*^+^] interval.

We performed meiotic mapping to ascertain the genetic basis of male sterility within each 2*P* introgression by generating recombinant 1*P* introgression genotypes (Figure 1B). 2*P*[*w*^+^] females were crossed to *D. simulans* strains carrying an X-linked *pBac*[*eYFP*] transgene (Stern et al., 2017) that served as an additional visible marker. Progeny from this cross were scored for recombinant X chromosomes carrying both *pBac*[*eYFP*] and a single *P*[*w*^+^] (1*P-YFP*). Recombinant 1*P-YFP* chromosomes were generated using *pBac*[*eYFP*] markers both proximal and distal to each 2*P* introgression. Virgin 1*P-YFP* females were individually crossed to *D. simulans w*^XD1^ males to initiate 1*P-YFP* strains. Each 1*P-YFP* X chromosome was then assayed for male fertility. At least 10 individual 1*P-YFP* males of each genotype were collected 1-2 days post-eclosion and aged 3-5 days, then placed singly in a vial with three virgin *D. simulans w*^XD1^ females. After seven days, both the male and females were discarded, and all offspring emerging from the vial were counted. Additional 1*P-YFP* males were archived for DNA extraction.

Progeny sex ratios were calculated as the number of female offspring/total number of offspring (% female). Males that sired fewer than five offspring were excluded from sex ratio analyses, as were genotypes with fewer than three males that sired more than four offspring. This resulted in 2538 males and 303 recombinant 1*P-*YFP chromosomes that were used to estimate progeny sex ratios; 210 recombinant 1*P-*YFP genotypes had both progeny sex ratio and sequence data.

### Genotyping recombinant chromosomes by sequencing

We determined the fine-scale genetic architecture of hybrid male sterility within each introgressed region by genotyping recombinant 1*P-YFP* X chromosomes using multiplexed whole-genome sequencing. DNA extraction and library construction followed published methods for high-throughput sequence analysis of a large number of recombinant genotypes (Andolfatto et al., 2011, Peluffo et al., 2015). Sequence reads were mapped to the reference genome sequence of the *D. mauritiana* stock used for mapping (*mau w*^12^) (Garrigan et al., 2012), our unpublished genome sequence of *sim w*^XD1^, and the *D. simulans pBac*[*eYFP*] strains (Stern et al., 2017). Ancestry from each parent species was determined by a Hidden Markov Model (HMM) (Andolfatto et al., 2011).

Across the 439 genotypes with sufficiently high-quality sequence data for ancestry assignment, we recovered 64,373 X-linked markers. A subset of 2,835 non-redundant markers were retained that delimit the extent of each 1*P-YFP D. mauritiana* segment. No genotype showed evidence for any autosomal *D. mauritiana* alleles (see Figure S2 for exemplars), confirming that our introgression scheme isolated X-linked *D. mauritiana* segments in a pure *D. simulans* autosomal genome.

### Samples and short read alignment

We used genome sequence data from 10 lines of *D. mauritiana*, including nine inbred wild isolates and the genome reference strain, *mau w*^12^(14021-0241.60); 20 lines of *D. simulans*, including 10 inbred wild isolates from Kenya (14021-0251.302-311), 9 wild isolates from Madagascar (14021-0251.293-301), and the reference strain, *sim w*^501^ (14021-0251.011); and the reference strain of *D. melanogaster*. The *D. mauritiana* and *D. simulans* sequence data were reported previously (Garrigan et al., 2012, Garrigan et al., 2014, Rogers et al., 2014). We performed short read alignment against the *D. mauritiana* genome assembly (version 2) using the “aln/sampe” functions of the BWA short read aligner and default settings (Li and Durbin, 2009). Reads flanking indels were realigned using the SAMTOOLS software (Li et al., 2009). Individual BAM files were merged and sorted with SAMTOOLS.

### Polymorphism and divergence analyses

Both within-and between-population summary statistics were estimated in 10-kb windows using the software package POPBAM (Garrigan, 2013). The within population summary statistics include: unbiased nucleotide diversity *π* (Nei, 1987); the summary of the folded site frequency spectrum Tajima’s *D* (Tajima, 1989); and the unweighted average pairwise value of the *r*^2^ measure of linkage disequilibrium, *ZnS*, excluding singletons (Kelly, 1997). The between population summary statistics include: two measures of nucleotide divergence between populations, *D*_*XY*_, and net divergence, *D*_*A*_ (Nei, 1987); the ratio of the minimum between-population nucleotide distance to the average, *G*_min_ (Geneva et al., 2015); and the fixation index, *F*_*ST*_ (Wright, 1951). In the analysis of absolute numbers of ancestral *versus* derived polymorphisms, we restricted the *D. simulans* sample to the 10 from Madagascar to enforce equal sample sizes with *D. mauritiana* (*n*=10). From a total of 11083 scanned 10-kb windows, we only analyzed windows for which at least 50% of aligned sites passed the default quality filters in POPBAM, which resulted in a final alignment for 10443 scanned 10-kb windows. POPBAM output was formatted for use in the R statistical computing environment using the package, POPBAMTools (https://github.com/geneva/POPBAMTools). All statistics and data visualization were done in R (Team, 2013).

### Identification of introgressed regions

We used the *G*_min_ statistic (Geneva et al., 2015) to scan the genome for haplotypes that have recent common ancestry between *D. simulans* and *D. mauritiana. G*_min_ is defined as the ratio of the minimum number of nucleotide differences per aligned site between sequences from different populations to the average number of nucleotide differences per aligned site between populations. The *G*_min_ statistic was calculated in 10-kb intervals across each major chromosome arm using the same quality filtering criteria used for all other summary statistics. From these values, we estimated the probability of the observed *G*_min_ under a model of allopatric divergence, conditioned on the divergence time. For each 10-kb interval, the significance of the observed *G*_min_ value was tested via Monte Carlo coalescent simulations of two populations diverging in allopatry with all mutations assumed to be neutral. We assumed a population divergence time of 1.21 × 2*N*_sim_ generations before the present, in which *N*_sim_ is the current estimated effective population size of *D. simulans* (Garrigan et al., 2012). In the simulations, the observed local value of *D*_*XY*_ was used to determine the neutral population mutation rate for that 10-kb interval. To account for uncertainty in local population recombination rate, for each simulated replicate, a rate was drawn from a normally distributed prior (truncated at zero) with the mean estimated from genetically determined crossover frequencies (True et al., 1996a). The empirical crossover rate estimates were converted from cM to *ρ* (the population crossover rate, 4*N*_sim_*c*, by assuming *N*_sim_ ≈10^6^). The effective population sizes of both species were assumed to be equal and constant. While the assumption of constant size is not realistic, it serves to make the test more conservative as any factor that decreases the effective population size also increases the rate of coalescence within populations, which has the net effect of increasing *G*_min_ and decreasing its variance, since fewer ancestral lineages will be available for inter-specific coalescence in the ancestral population (Geneva et al., 2015). For each 10-kb interval, 10^5^ simulated replicates were generated and the probability of the observed *G*_min_ value was estimated from the simulated cumulative density. To identify putatively introgressed haplotypes, we used a significance threshold of P ≤ 0.001 from the simulations, which yields a proportion of null tests of 0.982 and a false discovery rate of 5%. To infer the full length of any putative introgressions ≥ 10 kb, we identified runs of contiguous (or semi-contiguous) 10-kb windows with significant *G*_min_ values (*P* ≤ 0.001). Finally, we estimated maximum likelihood phylogenies for each of the putative introgression intervals using RAxML v. 8.1.1 (Stamatakis, 2014).

### Genotyping the Winters *sex ratio* genes

We extracted genomic DNA from single male flies using the Qiagen DNeasy Blood and Tissue Kit. The meiotic drive genes of the Winters *sex ratio* system (Tao et al., 2007a), *Dox* and *MDox*, were PCR-amplified as previously described (Kingan et al., 2010). To assay the presence or absence of the *Dox* and *MDox* gene insertions, the amplicons for the *Dox* and *MDox* regions were digested with the *StyI* and *StuI* restriction enzymes (NEB), respectively. The digests were run on a 1% agarose gel stained with EtBr and the band size was estimated using the GeneRuler 1 kb plus ladder (Thermo Scientific). For both genes, only haplotypes containing the gene insertions have restriction sites as confirmed by samples with known genotypes (Kingan et al., 2010).

### Quantitative PCR for *Dox/MDox* expression in fly testes

We assayed expression of the *Dox* and *MDox* genes in testes from *D. simulans* strain MD63 and *D. mauritiana* strain *mau w*^12^ using quantitative PCR. Total RNA was extracted from the dissected testes of 5-10 day old flies using the Nucleospin RNA XS kit (Macherey-Nagel, Germany), and cDNA was synthesized with poly dT oligos and random hexamers using Superscript III RT cDNA synthesis kit (Invitrogen, CA). qPCR assays were performed on a BioRad Real-time PCR machine using the cycling conditions: 95° C for 3 mins.; 40 cycles of 95° C for 10s, 58° C for 30s, and 72° C for 30s. The primer sequences used for qPCR are provided in **Table S1**.

## ACKNOWLEDGEMENTS

The authors thank Brian Calvi for generously providing space and Shelby Biel, Ally Shambaugh, and Amanda Meiklejohn for assistance with fertility assays.

---

**Figure S1.**
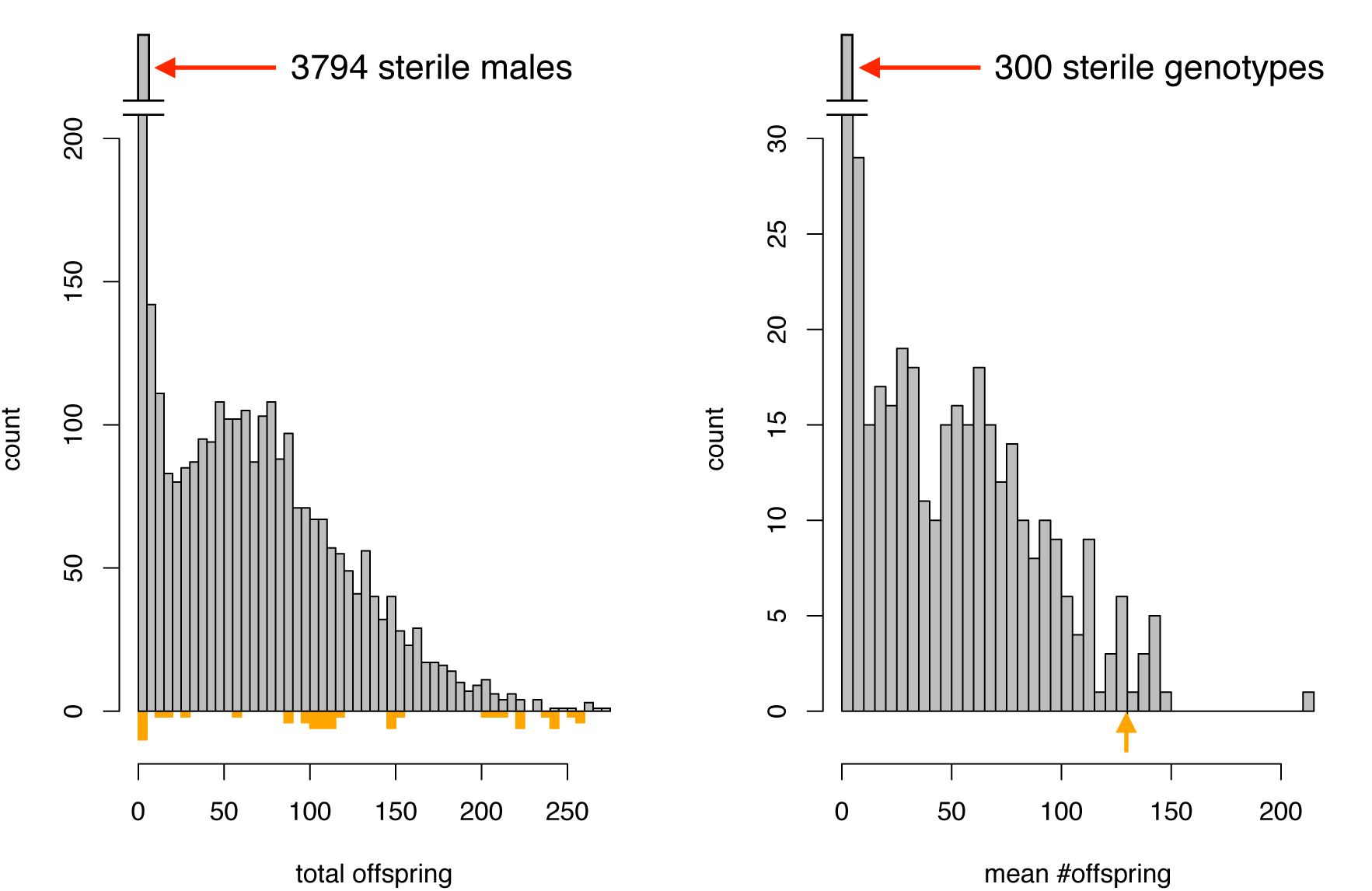
Distribution of fertility (number of progeny) among all males carrying recombinant 1*P-YFP* X chromosomes, and average number of progeny among all 1*P-YFP* genotypes. Colored bars and arrow below indicate individual male and mean fertility for *D. simulans w*^XD1^, respectively.

**Figure S2.**
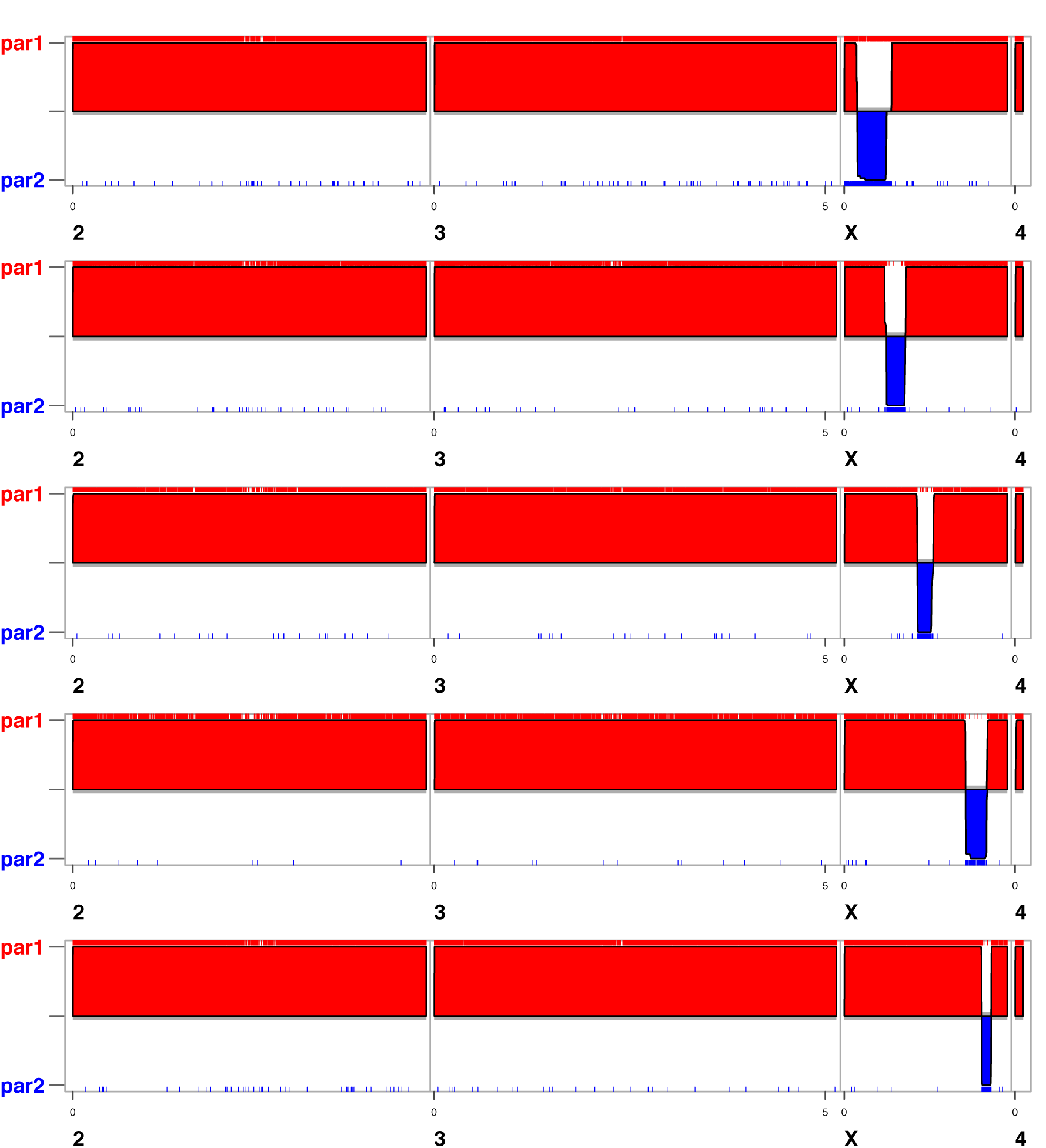
SNP locations and inferred ancestry for five recombinant 1*P-YFP* genotypes. Red ticks indicate *D. simuhms* alleles (pari), blue ticks indicate *D. mauriticma* alleles (par2). and the red (blue) shaded regions indicate the location of inferred *D. simulans (D. mauriticma*) ancestry.

**Figure S3.**
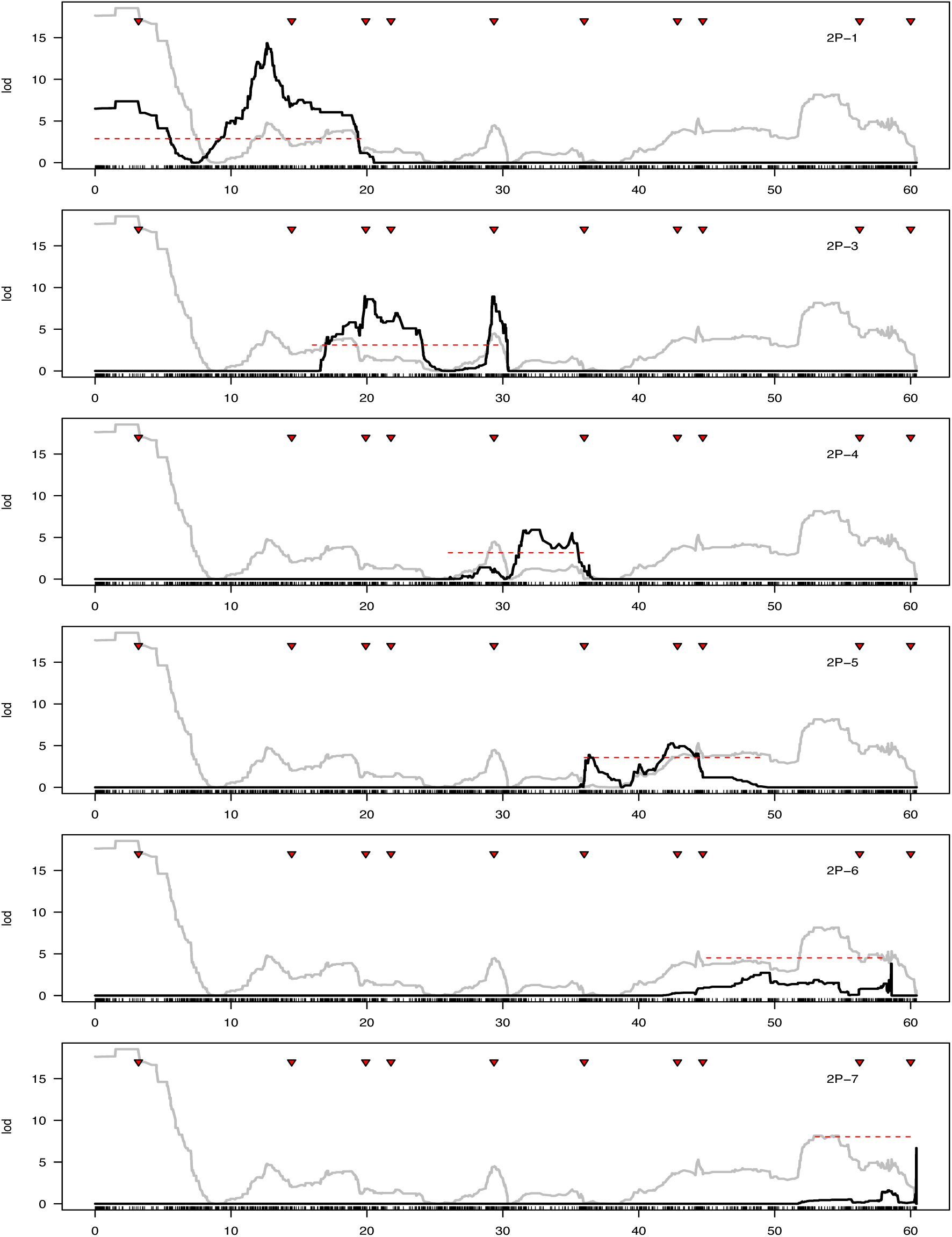
QTL analysis of male fertility. Recombinant 1*P-YFP* genotypes from each 2*P* introgression were used as separate mapping populations. Light grey lines in the background show LOD scores derived from using all 1*P-YFP* genotypes as a single mapping population (see Figure 3), red dashed line indicates significance threshold (*P* < 0.01) determined from 10.000 permutations.

**Figure S4.**
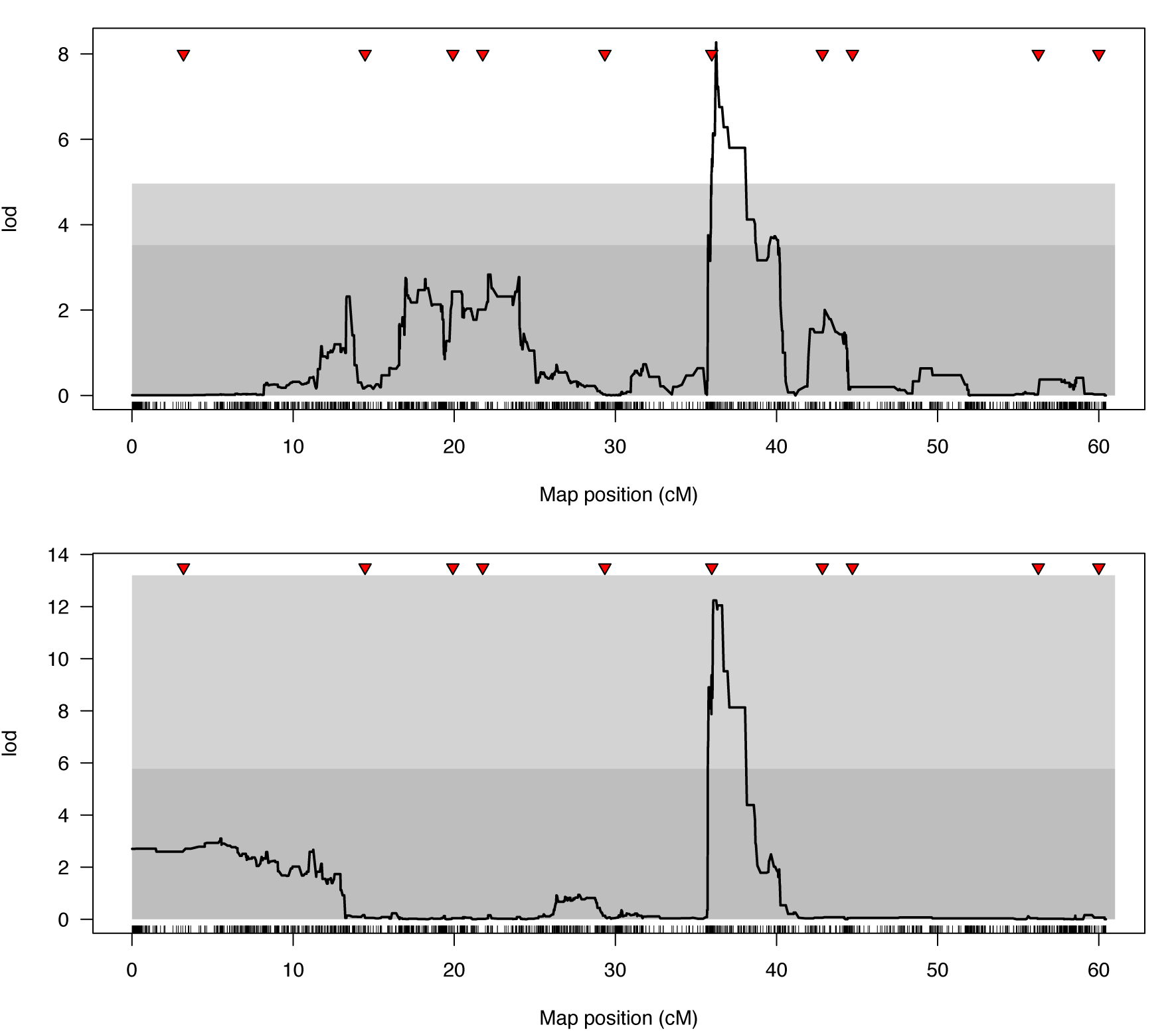
QTL analysis of progeny sex ratio associated with introgression genotypes. Dark grey and light grey regions indicate 5% and 1% significance thresholds determined from 10.000 random permutations. Top panel includes all males that produced any offspring; bottom panel includes only males that sired more than four offspring and genotypes with at least three males that sired more than four offspring.

